# Broad neutralizing nanobody against SARS-CoV-2 engineered from pre-designed synthetic library

**DOI:** 10.1101/2021.08.07.455523

**Authors:** Qianyun Liu, Chenguang Cai, Yanyan Huang, Li Zhou, Yanbin Guan, Shiying Fu, Youyou Lin, Ting Yang, Nanyan Wan, Fengzhi Zhang, Qi Sun, Ying Bai, Yu Chen, Xiaohua Liang, Huan Yan, Zhen Zhang, Ke Lan, Yu Chen, Xiang Li, Shin-Chen Hou, Yi Xiong

**Affiliations:** Bioduro-sundia LLC., Wuxi 214174, Jiangsu, China; State Key Laboratory of Virology, Modern Virology Research Center, College of Life Sciences, Wuhan University, Wuhan, 430072, Hubei, China

## Abstract

SARS-CoV-2 infection is initiated with Spike glycoprotein binding to the receptor of human angiotensin converting enzyme 2 via its receptor binding domain. Blocking this interaction is considered as an effective approach to inhibit virus infection. Here we report the discovery of a neutralizing nanobody, VHH60, directly produced from a humanized synthetic nanobody library. VHH60 competes with human ACE2 to bind the receptor binding domain of the Spike protein with a K_D_ of 2.56 nM, inhibits infections of both live SARS-CoV-2 and pseudotyped viruses harboring wildtype, escape mutations and prevailing variants at nanomolar level. VHH60 also suppresses SARS-CoV-2 infection and propagation 50-fold better and protects mice from death two times longer than that of control group after live virus inoculation on mice. VHH60 therefore is a powerful synthetic nanobody with a promising profile for disease control against COVID19.

## Introduction

The outbreak of coronavirus disease 2019 (COVID19) caused by a novel Severe Acute Respiratory Syndrome Coronavirus 2 (SARS -CoV-2) has lasted for more than two years with heavy tolls on human lives and deleterious consequences to international economic and social activities. Infected Individuals underwent estimated median viral incubation period of 7.8 days before showing clinical symptoms at 11.5 days post infection[1, 2]. Moreover, presymptomatic and around 15.6% asymptomatic patients both are contagious and contribute together more than 40% cases[3, 4]. These epidemic characteristics has seriously exposed the limitations of the existing epidemic prevention and control measures, thus it is urgent to develop efficacious vaccines and drugs to prevent virus transmission and antiviral therapy. SARS-CoV-2 is a single strand RNA virus that belongs to the genus Betacoronavirus, shares genomic identity of 79.6% to SARS-CoV and 96.2% to a bat coronavirus RaTG13[5-8]. Similar to other betacoronaviruses, SARS-CoV-2 infection is mediated by the viral glycoprotein Spike (S) through its binding to the host receptor Angiotensin Converting Enzyme 2 (ACE2). S protein is a trimeric fusion protein on the virion surface, which can be cleaved into receptor-binding fragment S1 and fusion fragment S2 by cellular serine protease TMPRSS2 and lysosomal proteases cathepsins upon engaging with the host cell [9, 10]. Another proprotein convertase Furin has also been reported to participate in SARS-CoV-2 entry by pre-activation of the S protein[10]. S1 firstly interacts with ACE2 through the receptor binding domain (RBD) on its C-terminus, then switches its conformation from “sitting-down” to “standing-up” to dissociate and expose S2 which functions mainly to drive virus fusion with the cell membrane. Although sharing the same receptor as SARS-CoV, SARS-CoV-2 S protein demonstrates stronger affinity with ACE2 due to different amino acids at the S1/S2 cleavage site[10, 11], which partially explains its higher transmissibility. The structure of the S protein, its binding pattern and its dynamic conformational changes during virus entry and fusion have been intensively analyzed at the atomic level[11-14], providing valuable guides to develop antiviral agents.

Since the first clinical trial of a vaccine against SARS-CoV-2 begun in March 2020[15], multiple vaccines with accelerated development have been deployed worldwide under Emergency Use Authorization (EUA). But efficacy and safety concerns were raised all the time[16, 17], reinfection after vaccination have been also reported[18, 19]. Nevertheless no drugs, such as monoclonal antibodies against SARS-CoV-2[20], have been formally approved for clinical use yet except EUA by the Food and Drug Administration (FDA). Among all monoclonal antibodies derived from human or small laboratory animals, a set of antibodies named nanobodies possess distinctive properties [21-23]. Nanobodies, also called VHH, are single domain antibody fragments originally found in llamas and other camelids. Unlike a conventional IgG antibody comprising covalently linked two light chains and heavy chains, a nanobody only has the monomeric target recognition module of the heavy chain but retains similar specificity and affinity. Given the small size of only 15 kD, nanobodies are very easy to produce and manipulate, featuring robust thermostability, solubility and permeability, yet low immunogenicity[24]. Nanobodies have been widely developed as therapeutic agents to treat cancer, autoimmune and renal diseases. Recently, a nanobody based drug has been successfully approved by FDA for clinical use, validating the single-domain construct as a new drug platform [25].

Recent S protein mutants such as E484K[26], N501Y[27], D614G[28, 29] and viral variants carrying more than one mutations like B1.1.7 (U.K. variant), B.1.351 (South African variant), P.1. (Brazilian variant), B.1.525(Nigeria variant) and B.1.617.2 (Delta variant) have been shown to gain enhanced binding to ACE2, viral replication and transmission, resulting in their capability to escape most of current neutralizing antibodies inhibition and vaccination prevention, with exacerbated pandemic and disease severity [26-28, 30-33]. Moreover, traditional approaches to generate and screen antibodies are usually initiated with animal immunization or with patient’s contagious samples. There are risks for researchers exposed to infectious pathogens and the immunization process also takes time. To overcome such scenario during global emergency, here we described a potent nanobody with broad neutralizing activity, which is synthesized and screened directly from pre-designed humanized nanobody library. Our data indicates this nanobody called VHH60 can bind to RBD with single digit nanomolar, blocking wildtype virus and variants entry, protect host cell and mouse from viral infection, suggests that VHH60 is a value candidate for further investigation to overcome COVID19. At another hand, our results also shed the light to discover therapeutical nanobodies safely and promptly.

## Material and Methods

### Cell lines and viruses

Vero-E6 (ATCC® CRL-1586), CaCO2(ATCC® HTB-37) and 293T (ATCC® CRL-3216) cell are cultured in Dulbecco’s Modified Eagle Medium (DMEM, Thermal Fisher, # 12430112), supplied with 10% fetal bovine serum (Thermal Fisher, # 26140079),1% antibiotic-antimyotic (Thermal Fisher, #15240096), and propagated at 37 °C in 5% CO2. ExpiCHO Expression System was purchased from Thermal Fisher (#A29133). SARS-CoV-2 Virus was provided by Dr. Yu Chen from Wuhan University.

### Protein expression and purification

The constructs of VHHs were selected from phage display, RBD fragment (aa319-541) of SARS-CoV-2 S protein (GenBank: MN908947.3) was synthesized by Genewiz Inc (GENEWIZ, Suzhou, China), the extracellular domain of human ACE2 (1-740 aa) (GenBank: NM_021804.1) was amplified from a plasmid (HG10108-ACG, Sinobiologic, Beijing, China). The whole coding cassette was ligated into a pCMV3 expression vector with a signal peptide of MEFGLSWVFLVALFRGVQC at the N-terminal, and either a 6-his tag (for RBD-his) or a human IgG1 Fc fragment with (GSSSS)3 linker at C-terminal. For Fc-tagged protein, the construct was expressed in 293F by transfecting with PEIMAX PEI MAX™ (24765-1, Polysciences, Inc., Warrington, PA, USA) according to the manual. The protein was purified from culture supernatant by protein-A affinity chromatography and was buffer changed into PBS and stored in -70 °C. For His-tagged protein, the construct was used for expression in CHO using ExpiCHO™ Expression according to the manual. The protein was purified from culture supernatant by Nickel affinity chromatography and finally dissolved in PBS (pH7.4) and stored in -70 °C. For VHH-Fc, the protein was purified from culture supernatant by protein A affinity chromatography and was buffer changed into PBS and stored in -70 °C. Extra polishing by a hydroxyapatite chromatography was added to achieve HPLC purity over 95% for specific experiments.

For protein A affinity chromatography, the culture supernatant from transient expression product was clarified by centrifuge at 3000g for 10min and was mixed with equal volume of 1.5M Glycine, 3M NaCl, pH8.9. the protein was purified with HiTrap™ MabSelect™SuRe™ column (11003494, Cytiva Inc., Marlborough MA, USA), and was eluted with 20mM acetic acid, pH3.5. The acid eluted fraction was neutralized with 1M Tris-HCl, pH9.0 and was concentrated and desalted into PBS with Amicon® Ultra-15, PLTK Ultracel-PL membrane (MilliporeSigma Life Science Center, Burlington, Massachusetts, USA) with appropriate MWCO.

For Nickel affinity chromatography, the culture supernatant from transient expression product was clarified by centrifuge at 3000g for 10min and was mixed with equal volume of 20mM imidazole, 500mM NaCl, 20mM Tris pH8.0. the protein was purified with HisTrap™ HP column (17524701, Cytiva Inc., Marlborough MA, USA), and was eluted with 500mM imidazole, 500mM NaCl, 20mM Tris pH8.0. The eluted fraction was concentrated and desalted into PBS with Amicon® Ultra-15 centrifugal unit (MilliporeSigma Life Science Center, Burlington, Massachusetts, USA) with appropriate MWCO. For polishing with hydroxyapatite chromatography, sample was buffer changed into 5mM sodium phosphate, 20mM MES, pH6.6, and was loaded on self-packed columns with Ca++Pure HA resin (45039, Tosoh Bioscience LLC, PA, Japan); the protein was eluted by a gradient elution with 400mM sodium phosphate, 20mM MES, pH6.6, and the targeted fraction was concentrated and desalted into PBS with Amicon® Ultra-15 centrifugal unit with appropriate MWCO. All proteins were checked by SDS-PAGE and HPLC with TSKgel G3000SWXL (08541, Tosoh Bioscience LLC, PA, Japan) for purity.

Construction of phage displayed synthetic VHH library by oligonucleotide-directed mutagenesis Three CDRs of the VHH template (a humanized VHH from the V germline gene, IGHV3S1*01, of Camelus dromedaries) were mutagenized using synthesized oligonucleotides coding tailored diversity of amino acids. In brief, mutagenic oligonucleotides for each CDR were mixed and phosphorylated by T4 polynucleotide kinase (New England BioLabs) in 70mM Tris–HCl (pH7.6), 10mM MgCl2, 1mM ATP and 5mM dithiothreitol (DTT) at 37□°C for 1h. The single stranded m13mp2 DNA template containing uracil was obtained from defective E. coli. The phosphorylated oligonucleotides were annealed to uracilated single-stranded DNA template at a molar ratio of 3:1 (oligonucleotide:ssDNA) by heating the mixture at 90□°C for 2□min, then followed by a temperature decrease of 1□°C/min to 20□°C in thermal cycler. Subsequently, the template-primer annealing mixture was incubated in 0.32mM ATP, 0.8mM dNTPs, 5mM DTT, 600□units of T4 DNA ligase and 75□units of T7 DNA polymerase (New England BioLabs) to prime in vitro DNA synthesis. After overnight incubation at 20□°C, the synthesized dsDNA was desalted and concentrated by a centrifugal filter (Amicon® Ultra 0.5mL 30K device), then electroporated into Escherichia coli ER2738 at 3000V followed by the M13KO7 helper phage infection and overnight culturing. Finally, the phage displaying nanobodies as a library in the culture medium was harvest and precipitated by polyethylene glycol (PEG)/NaCl for further use. Typically, 1μg of dU-ssDNA produced about 10^7^–10^8^ recombinant phage variants and 75–90% of the phage variants carried mutagenic oligonucleotides at three CDR regions simultaneously.

### Screening of anti-RBD nanobodies

RBD specific VHHs were identified from the screening (bio panning) of phage displayed synthetic VHH library. Recombinant RBD-Fc (2□∼□5μg per well) was coated in PBS buffer (pH7.4) in NUNC 96-well Maxisorb immunoplates overnight at 4°C and then blocked with 2% BSA in PBST for 1h. After blocking, 100 μL of resuspended polyethylene glycol (PEG)/NaCl-precipitated phage library (10^13^□cfu/mL in blocking buffer) was added to each well for 1h under gentle shaking. The plate was washed 10 times with 200 μL PBST [0.05% (v/v) Tween20] and 2 times with 200□μL PBS. The bound phages were eluted with 100 μL of 0.1M HCl/glycine (pH 2.2) per well, immediately neutralized with 8μL of 2M Tris-base buffer (pH9.1). The eluted phages were mixed with 1□mL of E. coli ER2738 (A600nm□=□0.6) for 30□min at 37□°C, ampicillin was added to eliminate uninfected bacteria. The bacterial culture was infected again with 100μL M13KO7 helper phage (∼10^11^□CFU total) at 37□°C for 1h and then added to 50mL of 2X YT medium containing kanamycin 50μg/mL and ampicillin 100μg/mL overnight at 37□°C with vigorous shaking. The rescued phage library was precipitated with 20% polyethylene glycol/NaCl and resuspended in PBS. The concentrated phage solution was used for the next round of panning. After 2–3 rounds of selection-amplification cycle, single colonies were randomly selected into deep 96-well culture plate containing 850 μL/well of 2YT (100 μg/mL ampicillin) and shook at 37°C for 3h. 50 μL M13KO7 (∼5□×□10^10^□CFU total) was then added to each well followed by adding 100 μL 2YT containing kanamycin (500 μg/mL) 1h later. The clones were incubated at 37□°C with vigorous shaking overnight then centrifuged at 3000g for 10 min at 4°C. 50 μL culture medium and 50μL 5% PBST milk was added to each well of three 96-well Maxisorb immunoplates pre-coated with antigen (1μg/ml) and blocked with 2% BSA in PBST. After 1h incubation at room temperature, the plates were washed six times with PBST. 100μL Protein M13-HRP antibody (1:3000) was added to each well for 1h. the plates were washed six times with PBST buffer and twice with PBS, developed for 3□min with 3,3’,5,5’-tetramethyl-benzidine peroxidase substrate (Kirkegaard & Perry Laboratories), quenched with 1.0M HCl and read spectrophotometrically at 450nm. Positive clones were selected by the following criteria: ELISA OD450□>□0.2 for the antigen-coated well (antigen binding positive); OD450□<□0.1 for negative control well. Unique clones were determined by sequencing the VHH DNA harbored in the phagemid.

### Screening of nanobodies blocking the interaction between RBD and hACE2

For blocking ELISA, a 96-well Maxisorp plate was coated with hACE2-Fc (2μg/ml, 100μl per well) at 4□°C, overnight and then blocked with blocking buffer (2% BSA in PBS) for 2□h. 50 ul of VHH-Fc cell supernatant (expression product of PCR fragment) was added to 50 ul of PBT, which containing RBD-his (40ng/ml). VHH72-Fc and a non-related VHH-Fc (produced with the same method) were used as a positive and negative control, respectively. hACE2-Fc (2ug/ml in PBS) was also as a reference. After 1□h incubation with gentle shaking, 90 ul of the mixtures were transferred to the BSA-blocked plate and incubated for 20 min. RBD-His binding to the plate was detected with anti-His tag mouse monoclonal antibody (1:3000 dilution, SinoBiological, 105327-MM02T) and followed by an HRP conjugated anti-mouse IgG (H+L) Goat antibody (Beyotime, A0216). The RBD binding signals were developed by 3,3’,5,5’-tetramethyl-benzidine peroxidase substrate (Kirkegaard & Perry Laboratories), quenched with 1.0□M HCl and read at OD 450□nm by a spectrophotometer.

### RBD binding assays

For ELISA-based binding assay, the RBD-Fc antigens (0.2µg per well) were coated in PBS buffer (pH7.4) on NUNC 96-well Maxisorb immunoplates overnight at 4 □, then blocked with 5% skim milk in PBST for 1h. 100µL nanobodies prepared at serial concentrations in PBST with 2.5% milk were added to each well and incubated for 1h under gentle shaking. The plate was washed with PBST and then added with 100µL 1: 2000-diluted anti-human IgG conjugated with horse-radish peroxidase for another 1h. The plates were washed with PBST buffer and twice with PBS, developed for 3min with 3,3’,5,5’-tetramethyl-benzidine peroxidase substrate (Kirkegaard & Perry Laboratories), quenched with 1.0 M HCl and read spectrophotometrically at 450nm. The EC50 was calculated according to Stewart and Watson method.

For HTRF based binding assay, serially diluted nanobodies were mixed with 50 nM biotinylated RBD for 15min at room temperature, then the 6μl mixtures were added to 4μl 10 nM hACE2-Fc diluted in 384-well plate for additional 15min. 10μl of detection mixture containing Streptavidin-d2 (Cisbio 610SADLF, 1:200), PAb Anti Human IgG-Eu cryptate (Cisbio 61HFCKLA, 1:200) was finally added and incubate for 60 min. signal were recorded by Envision (PerkinElmer, Inc) in FRET mode at wavelength of 665 nm. All reagents were diluted in assay buffer (PBS pH74,10% BSA), and proceeded at room temperature.

### Surface plasmon resonance (SPR) assay

The affinity of anti-RBD nanobodies and RBD antigen was measured with Biacore 8K. A biosensor chip, Series S Sensor Chip Protein A (Cat. # 29127556, GE), was used to affinity capture a certain amount of Fc tagged nanobodies to be tested and then flow through a series of COVID-19 S.P. RBD (Cat. # 40592-V08B, SB) under a concentration gradient on the surface of the chip (dilution ratio: 2, conc. levels: at least 5 (excluding curves with irregularities or high background)). Biacore 8K (GE) was used to detect the reaction signal in real time to obtain the binding and dissociation curves.

To measure the competitive response of anti-RBD nanobodies and hACE2, Fc-tagged nanobodies were immobilized on chip, then flowed with a 50nM of RBD (Cat. # 40592-V08B, SB) and a 100nM of hACE2 (Cat. # 1010B-H08H, SB). the reaction signal in real time were detected to obtain the binding and dissociation curves (theoretical ACE2 Rmax > 220 RU and kinetically simulated ACE2 binding > 160 RU for all). The buffer used in the experiment is HBS-EP^+^ solution (pH7.4, Cat. # BR100669, GE). The data obtained in the experiment was fitted with Biacore Insight Evaluation Software v3.0, GE software with a (1:1) binding model to obtain the affinity value.

### Pseudovirus neutralization

Pseudovirus neutralization assay was measured by reduction of Luciferase activity as described[34]. Briefly, pseudovirus bearing wildtype SARS-CoV-2 S protein or mutants were produced by co-transfection with plasmids expressing corresponding protein and backbone plasmid pNL-4-3-Luc.-R-E or VSV-dG-Lu. Pseudovirus was harvested, filtered and stored in -80°C. The pseudoviral titer was measured as 50% itssue culture infectious dose(TCID50) by infection of BHK21 overepressing human ACE2 according to the Reed-Muench method. Before infection of CaCO2 cells, pseudovirus was incubated with serial diluted nanobodies for 30min at room temperature. Luciferase activity was measured after 72 hours post infection according to manual of Bright-Glo™ Luciferase Assay System. Non-infected cells were considered as 100% inhibition, and cells only infected with virus were set as 0% inhibition. The IC50 values were calculated with non-linear regression using GraphPad Prism 8 (GraphPad Software, Inc., San Diego, CA, USA).

### SARS-CoV-2 neutralization assay

The SARS-CoV-2 (strain IVCAS 6.7512) was provided by the National Virus Resource, Wuhan Institute of Virology, Chinese Academy of Sciences. All SARS-CoV-2 live virus related experiments were approved by the Biosafety Committee Level 3 (ABSL-3) of Wuhan University. All experiments involving SARS-CoV-2 were performed in the BSL-3 and ABSL-3 facilities. Briefly, nanobodies were serially diluted in culture medium and 100□μl was mixed with 100□μl (1000 PFU) SARS-CoV-2 for 30 min.

The mixture was then added to Vero E6 cells in 48-well plates and incubated for 24□hours, after which TRIzol (Invitrogen) was added to inactivate SARS-CoV-2 viruses and extract RNA according to the manufacturer’s instructions. First-strand cDNA was synthesized using the PrimeScript RT kit (TakaRa). A real time quantitative PCR was used to detect the presence of SARS-CoV-2 viruses by the primers (table 1).

**Table 1:**
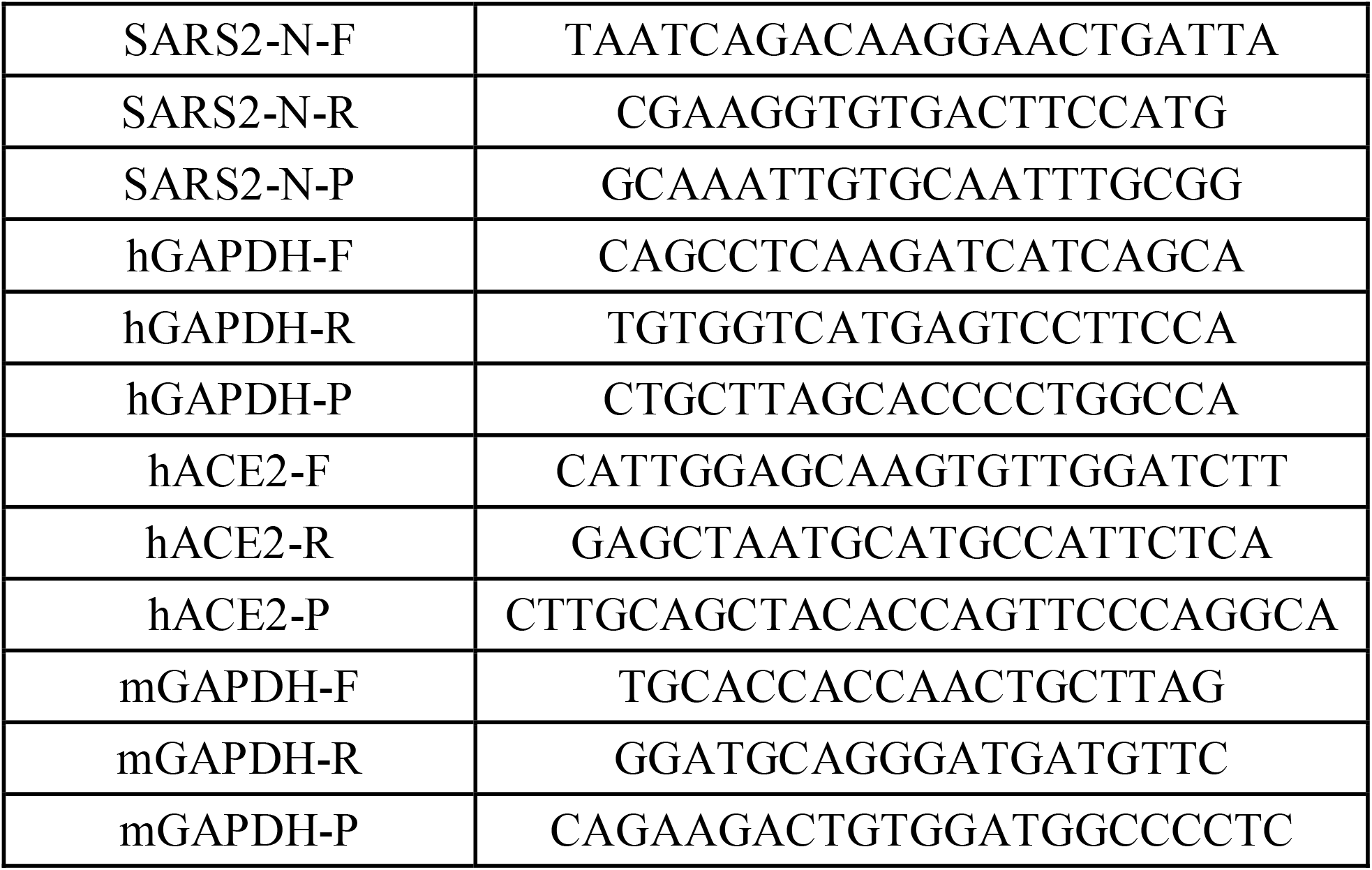
Primers:

The relative number of SARS-CoV-2 viral genome copies was determined using a TaqMan RT-PCR Kit (Yeason). To accurately quantify the absolute number of SARS-CoV-2 genomes, a standard curve was prepared by measuring the SARS-CoV-2 N gene constructed in the pCMV-N plasmid. All SARS-CoV-2 genome copy numbers were normalized to GAPDH expression in the same cell.

### Protection of K18-hACE2 transgenic mice against SARS-CoV-2

K18-hACE2 transgenic mice expressing human ACE2 driven by the human epithelial cell cytokeratin-18 (K18) promoter, were purchased from Gempharmatech and housed in ABSL-3 pathogen-free facilities under 12-h light-dark cycles with ad libitum access to food and water. All animal experiments were approved by the Animal Care and Use Committee of Wuhan University. Age-matched (9-10-week-old) female mice were grouped for infection of nanobodies (0.5mg/kg). One day later, mice were inoculated with 6 × 10^4^ PFU of SARS-CoV-2 by the intranasal route. Body weights were monitored at 3 and 6dpi.

Animals were sacrificed at 3 or 6 dpi according to the protocol, and tissues were harvested for pathologic and histologic analysis.

### Plaque assay of lung tissue homogenates

The right lung was homogenized in 1 mL PBS using a Tissue Cell-destroyer 1000 (NZK LTD). Vero E6 (ATCC number: CRL-1586) cells were cultured to determinate viral titer. Briefly, serial 10-fold dilutions of samples were added into monolayer cells. After adsorption at 37 □, the virus inoculum was removed and cells were washed with PBS twice, then DMEM containing 5 % FBS and 1.0 % methylcellulose was supplemented. Plates were incubated for 2 days until obvious plaques can be observed. Cells were stained with 1% crystal violet for 4h at room temperature. Plaques were counted and viral titers were defined as PFU/ml.

### Histological analysis

Lung samples were fixed with 4% paraformaldehyde, paraffin embedded and cut into 3.5-mm sections. Fixed tissue samples were used for hematoxylin-eosin (H&E) staining and indirect immunofluorescence assays (IFA). Wuhan Servicebio Technology Co., Ltd provides the histological analysis service. For IFA, Anti-hACE2 antibody and anti-SARS-CoV/SARS-CoV-2 Nucleocapsid Antibody (Cat: 10108-RP01 and 40143-MM05, SinoBiological) were added as primary antibodies. The image information was collected using a Pannoramic MIDI system (3DHISTECH, Budapest) and FV1200 confocal microscopy (Olympus).

### Data analysis

All the assays were conducted with at least duplicated biologic repeats. Results were presented by representative data or mean±SEM with indicated numbers of replication. All data was analyzed by XLfit (IDBS, Boston, MA 02210) or Prism 5(GraphPad Software, San Diego, CA 92108).

## Results

### Generation of neutralizing nanobodies against SARS-CoV-2

To generate the nanobodies that neutralize SARS-CoV-2, we used the pre-designed synthetic nanobody library technology. The complementary determination region (CDR) sequences with tailored diversity were genetically engineered into a humanized nanobody framework by a high-speed DNA mutagenesis method. The resulting nanobodies were displayed on phage surface as a library to be screened for binders against the recombinant RBD domain of the S protein. The phage displayed synthetic nanobody library of the size of 10^10^ were first screened against the RBD domain. Following 3 rounds of phage selections, the RBD specific clones were identified by single clone phage ELISA with positive signals (fig.1A). After sequencing, 78 unique VHH genes were identified and subsequently amplified by PCR from phage. Next, the mammalian expression cassette was assembled with a CMV promoter and a Fc domain of human IgG1 fragment on the VHH’s 5’ and 3’ end respectively by a second-round overlapping PCR. The final PCR products were transfected into ExpiCHO cells for Fc-tagged nanobody expression. Then the cell culturing medium containing Fc-tagged nanobodies was used to compete with Fc-tagged human ACE2(hACE2) for binding to the RBD as the 2^nd^ round ELISA-based blocking screening (fig.1B). 7 nanobodies were finally identified displaying blocking ability similar to that of the positive control VHH72-Fc[35]. All 7 antibodies were expressed for further characterization. As shown after protein A column purification, Fc tagged nanobodies were present as monomers of ∼40 kD in reducing gels with estimated purity of 90% (fig S1).

### Synthetic nanobodies specifically bind to SARS-CoV-2 RBD with high affinity

We next repeated ELISA to evaluate the binding capacity of the nanobodies to their original target RBD protein in a serial dilution manner. VHH35, VHH60, VHH79 and VHH80 showed slightly higher or comparable affinity against recombinant RBD protein compared to hACE2-Fc (fig.1C). Of all identified nanobodies, VHH35 displayed the highest affinity with an EC_50_ of 0.23 nM. Then we analyzed the affinity by surface plasmon resonance (SPR). VHH35 consistently exhibited the lowest K_D_ of 0.535 nM for RBD, the rest of nanobodies all bound to RBD with single digit nanomolar K_D_ values (fig. 1D, E). To confirm the blocking effect of nanobodies to the interaction between RBD and hACE2, we also used SPR to measure the binding of hACE2 to RBD after the RBD was pre-occupied by nanobodies captured by protein A on the chip. Our data indicated that after RBD was bound by all the tested nanobodies (including VHH34, VHH35, VHH43, VHH60, VHH79, VHH80, VHH82), no obvious binding curve of hACE2 to RBD could be detected, further demonstrating that these nanobodies have blocking effects to the binding of RBD to hACE2 (fig. S2). Together, the ELISA and SPR data indicated that these 7 nanobodies from the blocking screening have the ability to competing the binding of hACE2 to RBD.

**Figure 1.**
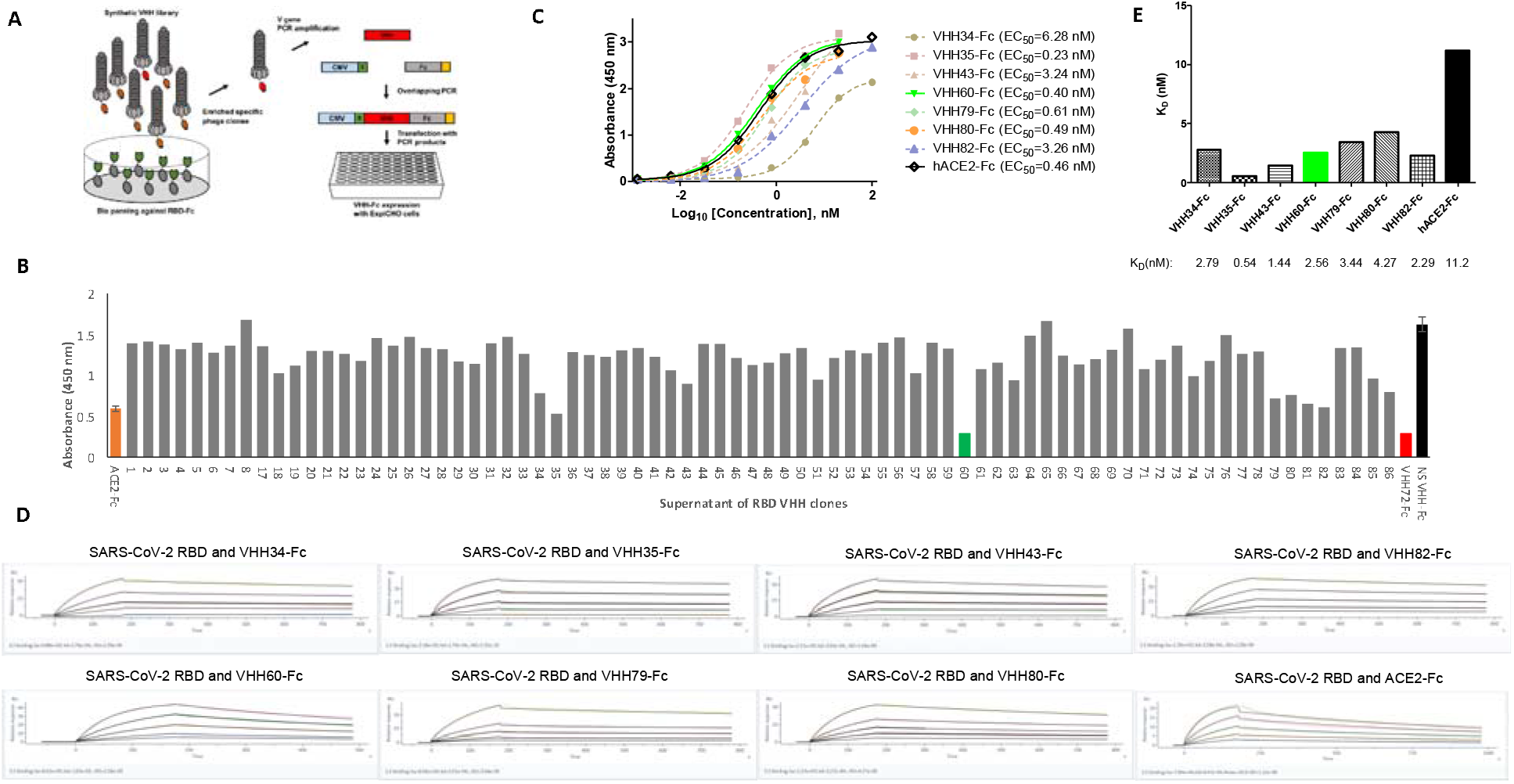
Screen and identify RBD specific nanobodies. A, the scheme of screening. B, Blocking ELISA to select nanobodies that can block hACE2 binding to RBD, smaller value reflects stronger blockage. Orange bar: hACE2-Fc as an internal control; Red bar: VHH72 as a positive control; Black bar: Non-reactive nanobody as a negative control. C, ELISA based binding assay. D, SPR. highest concentration: VHH34 (100 nM), VHH35 (50 nM), VHH43 (50 nM), VHH60 (25 nM), VHH79 (100 nM), VHH80 (100 nM), VHH82 (100 nM), hACE2-Fc (200 nM). E, chart of SPR results from E.

### VHH60 suppresses SARS-CoV-2 virus infection and amplification *in vitro* and *in vivo*

We then proceeded to investigate the neutralizing activity of VHH35 and VHH60 by pseudovirus based cell entry assay. Pseudovirus bearing the S protein and a luciferase was incubated with various concentrations of nanobodies for 30 min prior to infecting the CaCO2 cell. The resultant luciferase activity measured at 48 hours post infection suggested that VHH60 offered the best protection compared to VHH72 and ACE2, which are reflected by IC_50_ values of 13.96±2.42 nM, 26.34±5.19 nM and 16.81±5.06 nM respectively (fig.2A). Surprisingly VHH35 did not show expected efficacy with an IC_50_ value of around 100 nM (fig S3.) This is a case where binding affinity was not absolutely correlated with antiviral activity as observed by others [36].

To further evaluate the antiviral effect of VHH60, we used Authentical SARS-CoV-2 virus to test on Vero-E6 cell *in vitro*. Virus was premixed with serially diluted nanobodies for 30 min, then added to the Vero-E6 cell to propagate for 24 hours. Viral RNA level was measured by RT-PCR. The data showed that VHH60 inhibited viral infection at an IC_50_ of 1.87±0.35 nM, which was 6-fold lower than the IC_50_ of the reference VHH72 (11.17±1. 58 nM) (fig. 2B). Encouraged by this result, we then investigated the antiviral potential of VHH60 *in vivo*. 10 female K18-hACE2 transgenic mice per group expressing human ACE2 were intraperitoneally administrated with nanobodies or controls (Vehicle: PBS) at 0.5 mg/kg 24 hours prior to inoculation with authentical SARS-CoV-2 virus intranasally. 5 mice of each group were sacrificed for pathologic analysis at 3 d.p.i. as planned (fig. 2C). The remaining mice of the vehicle group all died at 4 d.p.i. (5 out of 5, observed at day 5), but mice treated with the nanobodies (VHH60 and VHH72) survived up to 6 days excepted one mouse in the VHH60 group that died at 5 d.p.i. (1 out of 5, observed at day 6) which could be considered as a normal variation (fig. 2D). All VHH60 and VHH72 treated mice were sacrificed at day 6 post infection because the body weight of the mice dropped up to 25%, which met the termination criteria according to the IACUC protocol. Consistent with previous report that virus infection could cause body weight loss [37], we also observed that at 3 d.p.i. the body weight of mice from the vehicle group had dropped 20%. In contrast the body weight of mice treated with VHH60 and VHH70 decreased only slightly (fig. 2E). To more accurately assess the protective effect, viral load was evaluated at 3 d.p.i when all mice including the vehicle group were still alive. Virus titer from lung in VHH60 treated group was significantly suppressed to a level which was 45-fold lower than that of the vehicle and 9-fold of the VHH72 group, respectively (fig. 2F). Immunofluorescent data clearly confirmed that the viral particles represented by green nucleocapsid staining were much fewer in VHH60 and VHH72 groups compared to those in the vehicle group (fig. 2G). We did not observe any significant difference of red signal from ACE2 staining which could exclude the possibility that virus titer was affected by the ACE2 level. Together, the results strongly support that VHH60 is highly efficacious to restrain SARS-CoV-2 infection and proliferation both *in vitro* and *in vivo*, ameliorate disease progress and improve health.

**Figure 2.**
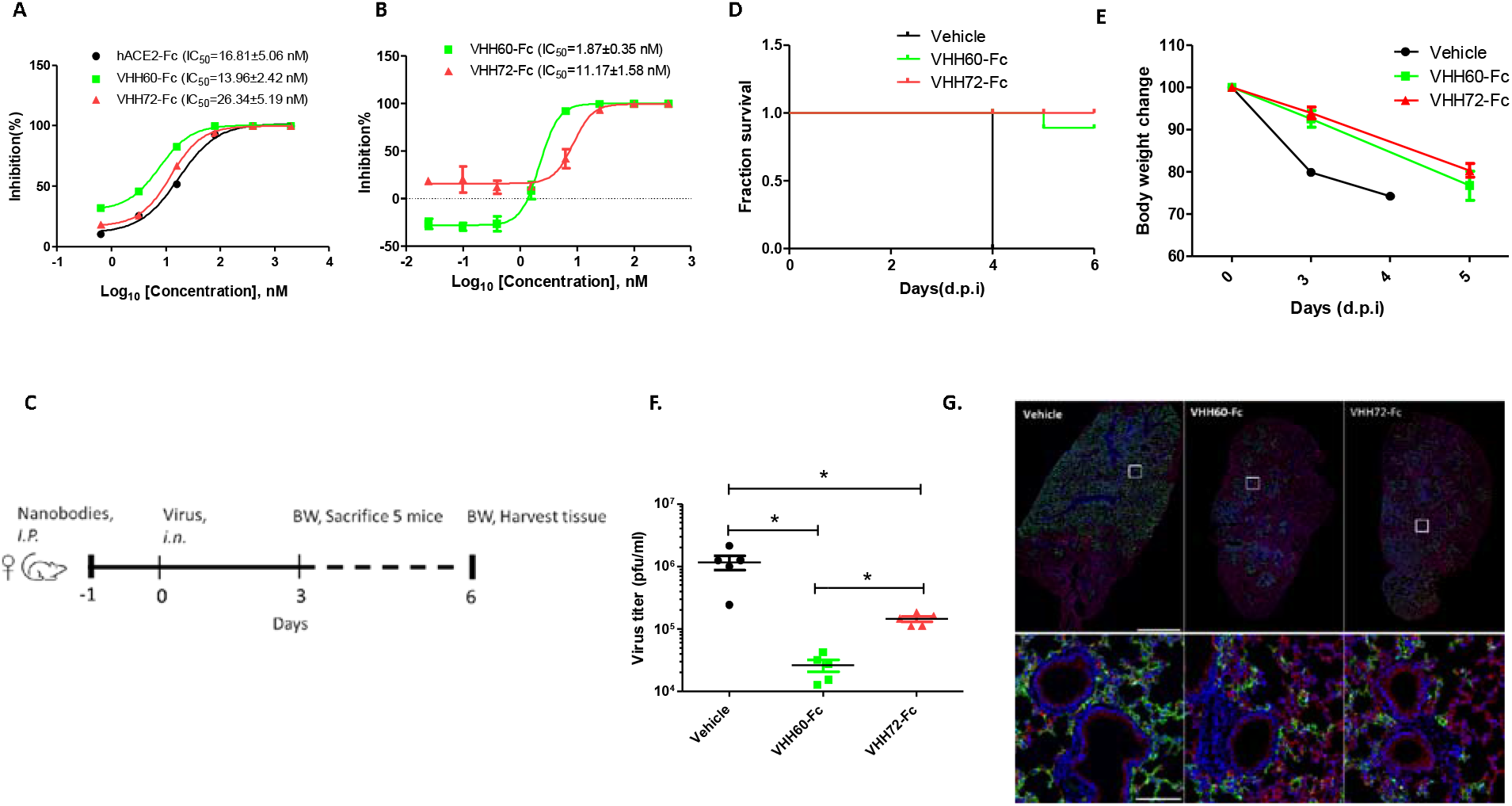
Antiviral activity of VHH60. A, VHH60 inhibits pseudovirus carrying wildtype spike protein infection on CaCO2 cell line (n=4). B, VHH60 inhibits authentic SARS-CoV-2 infection on Vero-E6 (n=2). C, Scheme of animal challenge. Total 10 mice were separated in each group, 5 mice were sacrificed 3 d.p.i. all remaining mice will be terminated after meeting certain criteria. D, Survival curve of mice infected by authentic SARS-CoV-2. Vehicle group all died at 4 d.p.i (5/5), one in VHH60 group (1/5) died. E, Body weight change of mice infected by authentic SARS-CoV-2. Data is represented as ratio of body weights at indicated timepoint versus day 0 (n=10 at 0 and 3 d.p.i, n=4 at 4 d.p.i, n=5 at 5.d.p.i). F, Virus load in lung after 3 days of infection (n=5). G, Representative image of immunofluorescence from lung. Blue: Nuclei, Red: ACE2, Green: nucleocapsid protein (NP). Upper panel: whole section, Lower panel: zoomed from white squares at upper panel (n=3). *:P<0.05

### VHH60 blocks the infection of mutated pseudovirus

Given the high mutagenic capacity of SARS-CoV-2 as an RNA virus, escape mutations and variants resistant to current antibodies or vaccines have been studied and described [26-33]. Antibody cocktails or broadly neutralizing antibodies have stood out and gained more attentions as a new modality to combat COVID19[38, 39]. In addition to the wildtype S protein, mutants and variants carrying more than one mutation were tested with VHH60 in pseudoviral entry assays. VHH60 inhibited all single mutants E484K, N501Y and D614 at nanomolar IC_50_ level (fig. 3A). Strikingly, VHH60 also exhibited robust activity to suppress variants B1.1.7 (IC_50_ =31.76 nM), B.1.351(IC_50_ =18.28 nM), P.1 (IC_50_ = 16.29 nM), and B.1.525 (IC_50_ =15.0 nM) at IC_50_ values close to or even better than that for wildtype S protein (fig 3B).

**Figure 3.**
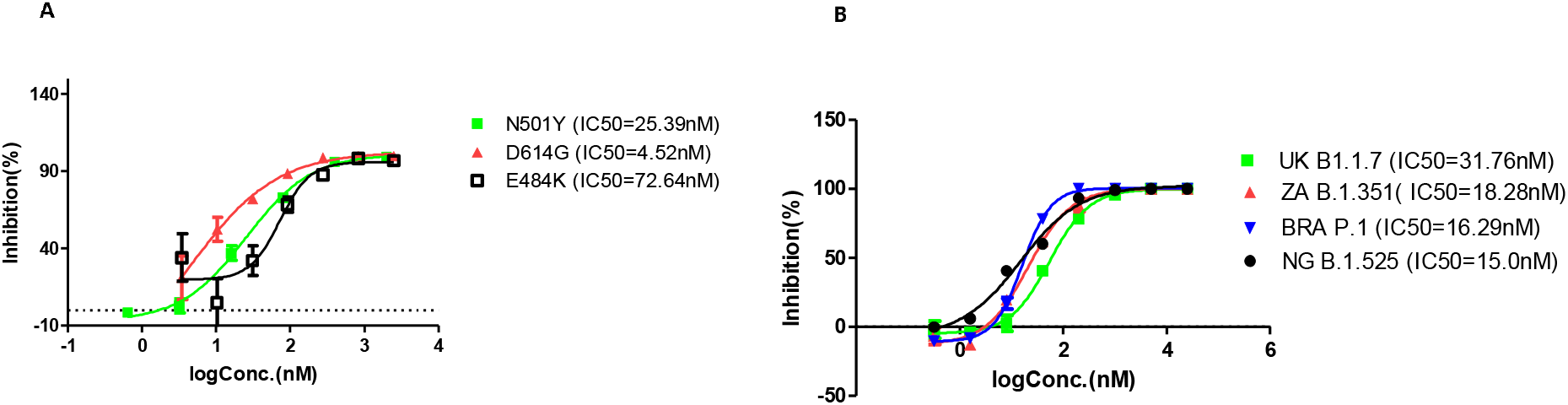
VHH60 blocks escape mutants and prevailing variants. A, VHH60 inhibits pseudovirus carrying spike protein with single mutation from infecting CaCO2 cell line (n=2). B, VHH60 inhibits pseudovirus carrying spike protein with multiple mutations as in reported variants from infecting CaCO2 cell line.

## Discussion

Overall, our results demonstrate that VHH60 potently competes with hACE2 to bind RBD and effectively blocks SARS-CoV-2 virus interaction with its host receptor to prevent infection in cell lines and a mouse model. Importantly VHH60 also maintains a broad capacity to neutralize multiple escape mutants and variants. Nanobodies have been considered as an important innovation for antibody-based therapies.

Unlike traditional monoclonal antibodies, nanobodies have the natural advantage for prophylactic applications, which are especially important considering SARS-CoV-2 is mainly spread via droplets and aerosol[40].

It is well known that multivalent nanobodies may have even higher affinity and antiviral activity, so we generated a His-tagged trimeric VHH60 and tested its antiviral activity. Compared to its monomeric form, the Tri-VHH60-His construct showed 9-times higher potency to block ACE2 interacting with RBD in a HTRF assay with an IC_50_ of 33.57 nM (fig. S4). Consequently, Tri-VHH60-His also provided superior inhibition of pseudovirus infection which is 9-times better than the monomeric VHH60 (fig. S5). Clearly, VHH60 can serve as a fertile basis for further optimizations.

During the COVID19 pandemic, more than 300 nanobodies from difference sources have been tested with antiviral activities, some of which showed up to femtomolar affinity to RBD in *in vitro* assays [37], suggesting that nanobodies could potentially be developed into versatile medications to combat COVID19. In this regard, the VHH60 nanobody reported here has demonstrated excellent potentials for further optimization and development towards clinical applications, based on its broad neutralizing capacity for both wildtype and most reported mutant forms of the SARS-CoV-2 virus. We have also shown that a pre-designed synthetic antibody library can be advantageous over the traditional antibody libraries, allowing us to rapidly discover functional synthetic antibodies without the lengthy process of animal immunization and hazardous exposure to pathogens. The synthetic nanobody platform described here can be readily deployed to discover antiviral molecules in the future when new viruses emerge.

## Acknowledgement

We thank Ming Guo, Xin Wang, Zhixiang Huang from ABLS3 laboratory for technical support. This work was supported by the Wuxi Municipal Bureau of Science and Technology (WX03-02B0136-092000-18) and grants from the National Science and Technology Major Project (2018YFA0900801), China NSFC projects (32041007), Fundamental Research Funds for the Central Universities (2042021kf0220), the Advanced Customer Cultivation Project of Wuhan National Biosafety Laboratory (2021ACCP-MS10) and Special Fund for COVID-19 Research of Wuhan University.

## Conflict of Interest Statement

Bioduro-sundia LLC. has a pending patent application for nanobodies listed in this publication except VHH72. There is no another competing interest.

## Notes

### Competing Interest Statement

The authors have declared no competing interest.

## References

1. Jing Qin, C.Y., Qiushi Lin Taojun, Shicheng Yu, Xiao-Hua Zhou, Estimation of incubation period distribution of COVID-19 using disease onset forward time: a novel cross-sectional and forward follow-up study. Science Advances, 2020. 6(33).

2. Lauer, S.A., et al., The Incubation Period of Coronavirus Disease 2019 (COVID-19) From Publicly Reported Confirmed Cases: Estimation and Application. Ann Intern Med, 2020. 172(9): p. 577–582.

3. He, X., et al., Temporal dynamics in viral shedding and transmissibility of COVID-19. Nat Med, 2020. 26(5): p. 672–675.

4. He, J., et al., Proportion of asymptomatic coronavirus disease 2019: A systematic review and meta-analysis. J Med Virol, 2020.

5. Chen, Y., Q. Liu, and D. Guo, Emerging coronaviruses: Genome structure, replication, and pathogenesis. J Med Virol, 2020.

6. Wu, F., et al., A new coronavirus associated with human respiratory disease in China. Nature, 2020. 579(7798): p. 265–269.

7. Lu, R., et al., Genomic characterisation and epidemiology of 2019 novel coronavirus: implications for virus origins and receptor binding. Lancet, 2020. 395(10224): p. 565–574.

8. Zhou, P., et al., A pneumonia outbreak associated with a new coronavirus of probable bat origin. Nature, 2020. 579(7798): p. 270–273.

9. Hoffmann, M., et al., SARS-CoV-2 Cell Entry Depends on ACE2 and TMPRSS2 and Is Blocked by a Clinically Proven Protease Inhibitor. Cell, 2020. 181(2): p. 271–280 e8.

10. Shang, J., et al., Cell entry mechanisms of SARS-CoV-2. Proc Natl Acad Sci U S A, 2020. 117(21): p. 11727–11734.

11. Shang, J., et al., Structural basis of receptor recognition by SARS-CoV-2. Nature, 2020. 581(7807): p. 221–224.

12. Wrapp, D., et al., Cryo-EM structure of the 2019-nCoV spike in the prefusion conformation. Science, 2020. 367(6483): p. 1260–1263.

13. Cai, Y., et al., Distinct conformational states of SARS-CoV-2 spike protein. Science, 2020. 369(6511): p. 1586–1592.

14. Yan, R., et al., Structural basis for the recognition of SARS-CoV-2 by full-length human ACE2. Science, 2020. 367(6485): p. 1444–1448.

15. Krammer, F., SARS-CoV-2 vaccines in development. Nature, 2020. 586(7830): p. 516–527.

16. Cines, D.B. and J.B. Bussel, SARS-CoV-2 Vaccine-Induced Immune Thrombotic Thrombocytopenia. N Engl J Med, 2021.

17. Tau, N., D. Yahav, and D. Shepshelovich, Vaccine safety - is the SARS-CoV-2 vaccine any different? Hum Vaccin Immunother, 2021. 17(5): p. 1322–1325.

18. Keehner, J., et al., SARS-CoV-2 Infection after Vaccination in Health Care Workers in California. N Engl J Med, 2021.

19. Teran RA W.K., Shane EL, et al., Postvaccination SARS-CoV-2 Infections Among Skilled Nursing Facility Residents and Staff Members — Chicago, Illinois, December 2020–March 2021. MMWR Morb Mortal Wkly Rep, 2021.

20. Anti-SARS-CoV-2 Monoclonal Antibodies. https://www.covid19treatmentguidelines.nih.gov/anti-sars-cov-2-antibody-products/anti-sars-cov-2-monoclonal-antibodies/, 2021.

21. Martinez-Delgado, G., Inhaled nanobodies against COVID-19. Nat Rev Immunol, 2020. 20(10): p. 593.

22. Schoof, M., et al., An ultra-high affinity synthetic nanobody blocks SARS-CoV-2 infection by locking Spike into an inactive conformation. bioRxiv, 2020.

23. Xiang, Y., et al., Versatile, Multivalent Nanobody Cocktails Efficiently Neutralize SARS-CoV-2. bioRxiv, 2020.

24. Muyldermans, S., Nanobodies: natural single-domain antibodies. Annu Rev Biochem, 2013. 82: p. 775–97.

25. Morrison, C., Nanobody approval gives domain antibodies a boost. Nat Rev Drug Discov, 2019. 18(7): p. 485–487.

26. Jangra, S., et al., SARS-CoV-2 spike E484K mutation reduces antibody neutralisation. Lancet Microbe, 2021.

27. Liu, H., et al., The basis of a more contagious 501Y. V1 variant of SARS-COV-2. Cell Research, 2021.

28. Zhou, B., et al., SARS-CoV-2 spike D614G change enhances replication and transmission. Nature, 2021. 592(7852): p. 122–127.

29. Hou, Y.J., et al., SARS-CoV-2 D614G variant exhibits efficient replication ex vivo and transmission in vivo. Science, 2020. 370(6523): p. 1464–1468.

30. Liu, Z., et al., Identification of SARS-CoV-2 spike mutations that attenuate monoclonal and serum antibody neutralization. Cell Host Microbe, 2021. 29(3): p. 477–488 e4.

31. Garcia-Beltran, W.F., et al., Multiple SARS-CoV-2 variants escape neutralization by vaccine-induced humoral immunity. Cell, 2021.

32. Starr, T.N., et al., Complete map of SARS-CoV-2 RBD mutations that escape the monoclonal antibody LY-CoV555 and its cocktail with LY-CoV016. Cell Rep Med, 2021. 2(4): p. 100255.

33. Hacisuleyman, E., et al., Vaccine Breakthrough Infections with SARS-CoV-2 Variants. N Engl J Med, 2021.

34. Nie, J., et al., Establishment and validation of a pseudovirus neutralization assay for SARS-CoV-2. Emerg Microbes Infect, 2020. 9(1): p. 680–686.

35. Wrapp, D., et al., Structural Basis for Potent Neutralization of Betacoronaviruses by Single-Domain Camelid Antibodies. Cell, 2020. 181(5): p. 1004–1015 e15.

36. Wec, A.Z., et al., Broad neutralization of SARS-related viruses by human monoclonal antibodies. Science, 2020. 369(6504): p. 731–736.

37. Rogers, T.F., et al., Isolation of potent SARS-CoV-2 neutralizing antibodies and protection from disease in a small animal model. Science, 2020. 369(6506): p. 956–963.

38. Xiang, Y., et al., Versatile and multivalent nanobodies efficiently neutralize SARS-CoV-2. Science, 2020.

39. Stamatatos, L., et al., mRNA vaccination boosts cross-variant neutralizing antibodies elicited by SARS-CoV-2 infection. Science, 2021.

40. Transmission of SARS-CoV-2: implications for infection prevention precautions. https://www.who.int/news-room/commentaries/detail/transmission-of-sars-cov-2-implications-for-infection-prevention-precautions, 2020.

