## Supplemental materials for "Broad neutralizing nanobody against SARS-CoV-2 engineered from pre-designed synthetic library"

**Figure S1.** Purified nanobodies with Fc tag were subjected to SDS-PAGE. Left panel: protein sample was reduced by DTT; Right panel: non-reducing samples. Each lane was labeled with the number corresponding to the nanobodies at far right.

**Figure S2.** Nanobodies inhibit interaction between RBD and hACE2 by SPR. The Fc tagged nanobodies and a reference antibody (Novoprotein Neutralizing Antibody) were captured onto the Protein A Chip as indicated at the first curve. The second binding curve was detected when 50 nM RBD (S.P.RBD) was injected. Lastly, injection of 100 nM hACE2 showed no further binding curve in all the experiments.

**Figure S3.** VHH35 inhibits pseudovirus carrying wildtype spike protein infection on CaCO2 cell line. The value of $IC_{50}$ is the average of two independent experiments.

**Figure S4.** Trimeric VHH60 Inhibits interaction of RBD and hACE2 by HTRF.

**Figure S5.** Trimeric VHH60 inhibits pseudovirus carrying wildtype spike protein infection on CaCO2 cell line. The value of $IC_{50}$ is the average of two independent experiments.

Figure. S1

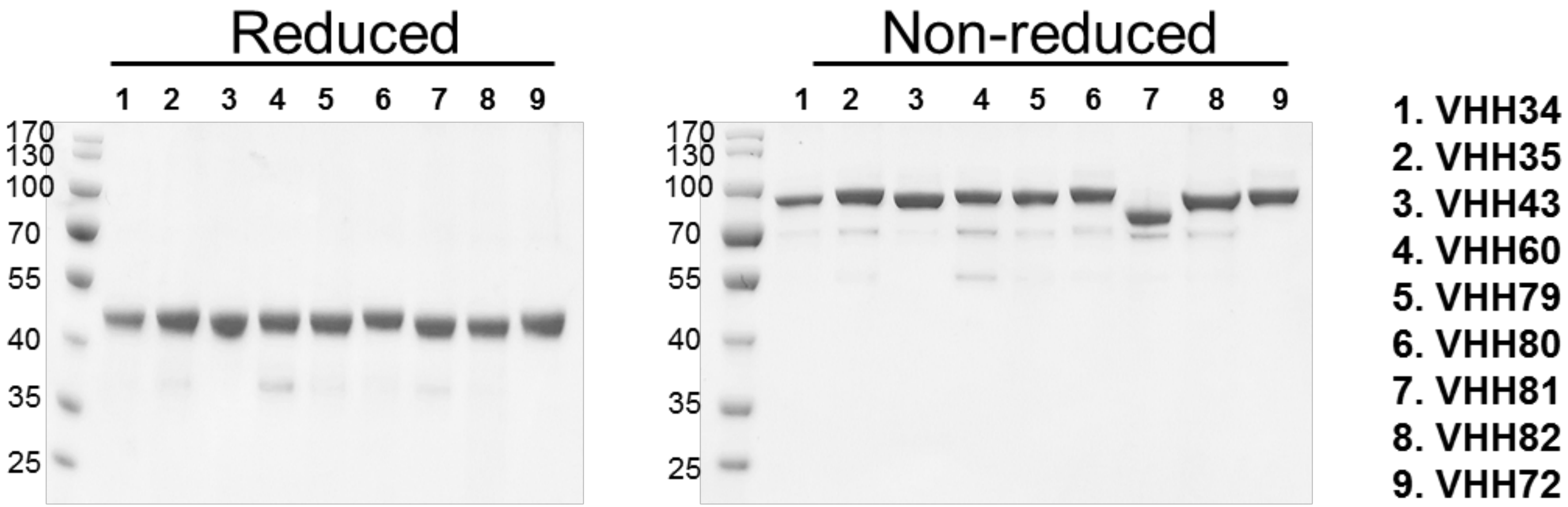
Reduced
1 2 3 4 5 6 7 8 9
170
130
100
70
55
40
35
25
Non-reduced
1 2 3 4 5 6 7 8 9
170
130
100
70
55
40
35
25
1. VHH34
2. VHH35
3. VHH43
4. VHH60
5. VHH79
6. VHH80
7. VHH81
8. VHH82
9. VHH72

### Figure.S2

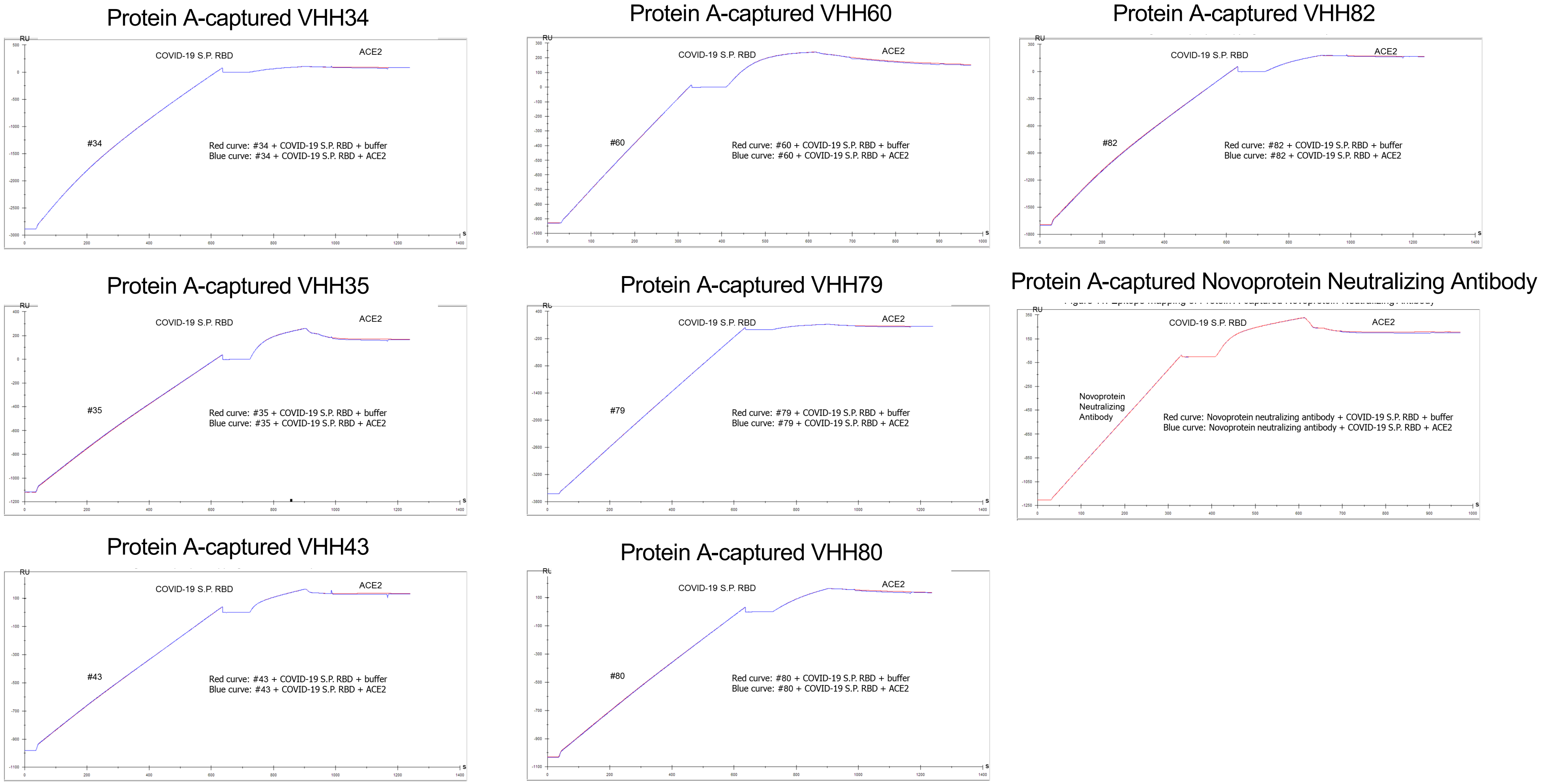

Protein A-captured VHH34
RU
COVID-19 S.P. RBD
ACE2
#34
Red curve: #34 + COVID-19 S.P. RBD + buffer
Blue curve: #34 + COVID-19 S.P. RBD + ACE2
s
Protein A-captured VHH60
RU
COVID-19 S.P. RBD
ACE2
#60
Red curve: #60 + COVID-19 S.P. RBD + buffer
Blue curve: #60 + COVID-19 S.P. RBD + ACE2
s
Protein A-captured VHH82
RU
COVID-19 S.P. RBD
ACE2
#82
Red curve: #82 + COVID-19 S.P. RBD + buffer
Blue curve: #82 + COVID-19 S.P. RBD + ACE2
s
Protein A-captured VHH35
RU
COVID-19 S.P. RBD
ACE2
#35
Red curve: #35 + COVID-19 S.P. RBD + buffer
Blue curve: #35 + COVID-19 S.P. RBD + ACE2
s
Protein A-captured VHH79
RU
COVID-19 S.P. RBD
ACE2
#79
Red curve: #79 + COVID-19 S.P. RBD + buffer
Blue curve: #79 + COVID-19 S.P. RBD + ACE2
s
Protein A-captured Novoprotein Neutralizing Antibody
RU
COVID-19 S.P. RBD
ACE2
Novoprotein
Neutralizing
Antibody
Red curve: Novoprotein neutralizing antibody + COVID-19 S.P. RBD + buffer
Blue curve: Novoprotein neutralizing antibody + COVID-19 S.P. RBD + ACE2
s
Protein A-captured VHH43
RU
COVID-19 S.P. RBD
ACE2
#43
Red curve: #43 + COVID-19 S.P. RBD + buffer
Blue curve: #43 + COVID-19 S.P. RBD + ACE2
s
Protein A-captured VHH80
RU
COVID-19 S.P. RBD
ACE2
#80
Red curve: #80 + COVID-19 S.P. RBD + buffer
Blue curve: #80 + COVID-19 S.P. RBD + ACE2
s

Figure. S3

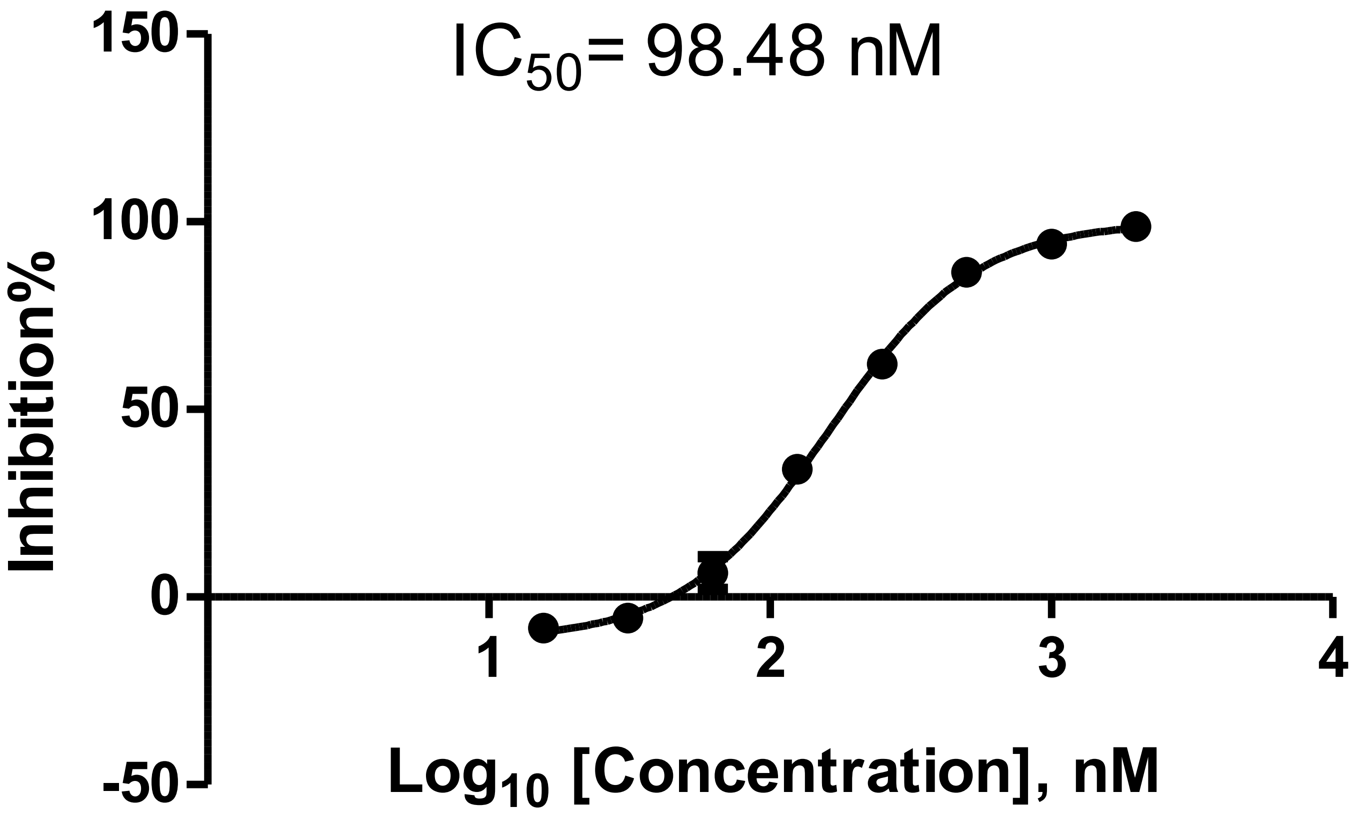

IC50= 98.48 nM
150
100
50
0
-50
Inhibition%
1
2
3
4
Log10 [Concentration], nM

### Figure. S4

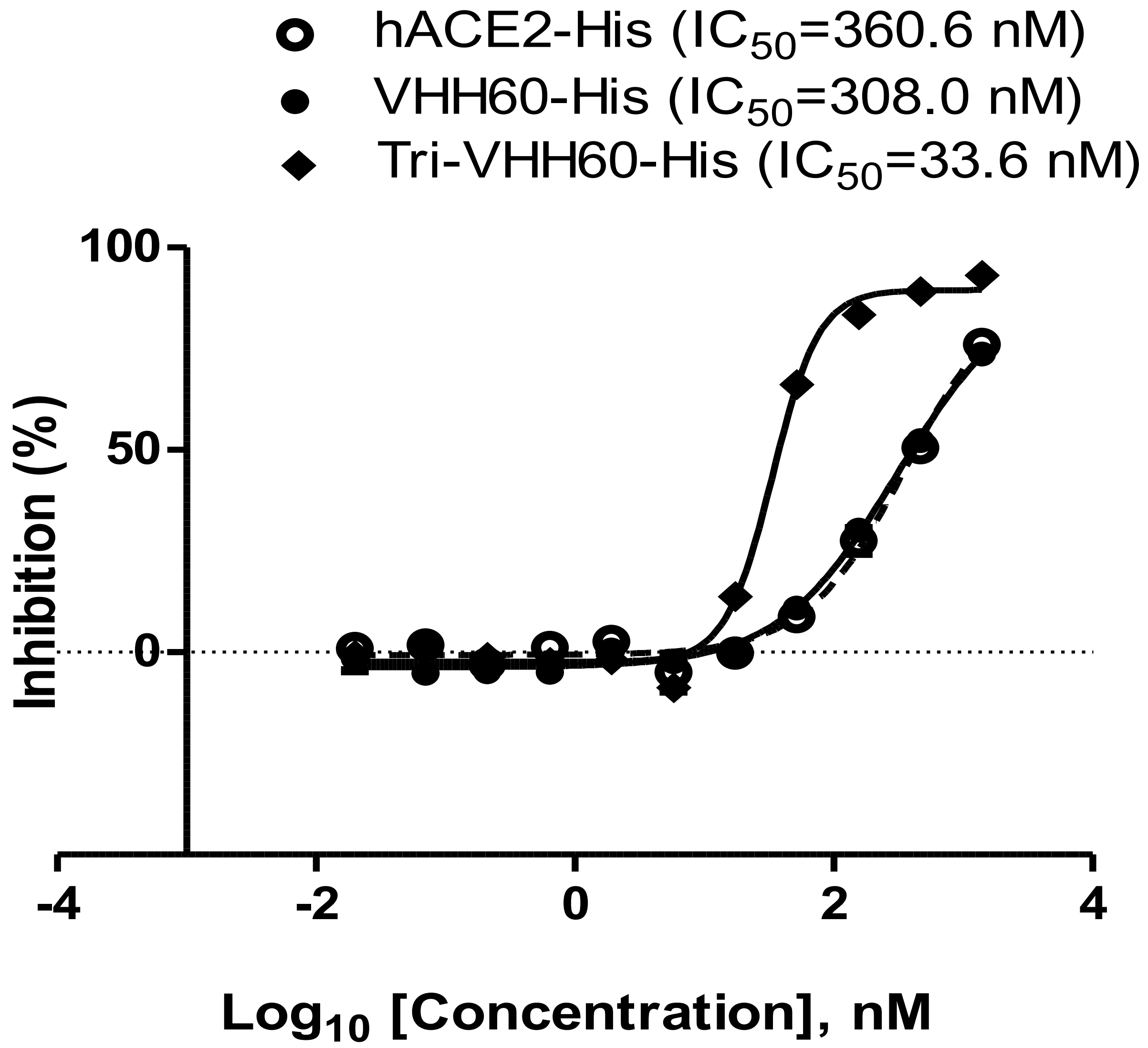
hACE2-His (IC50=360.6 nM)
VHH60-His (IC50=308.0 nM)
Tri-VHH60-His (IC50=33.6 nM)
100
50
0
Inhibition (%)
-4
-2
0
2
4
Log10 [Concentration], nM

Figure. S5

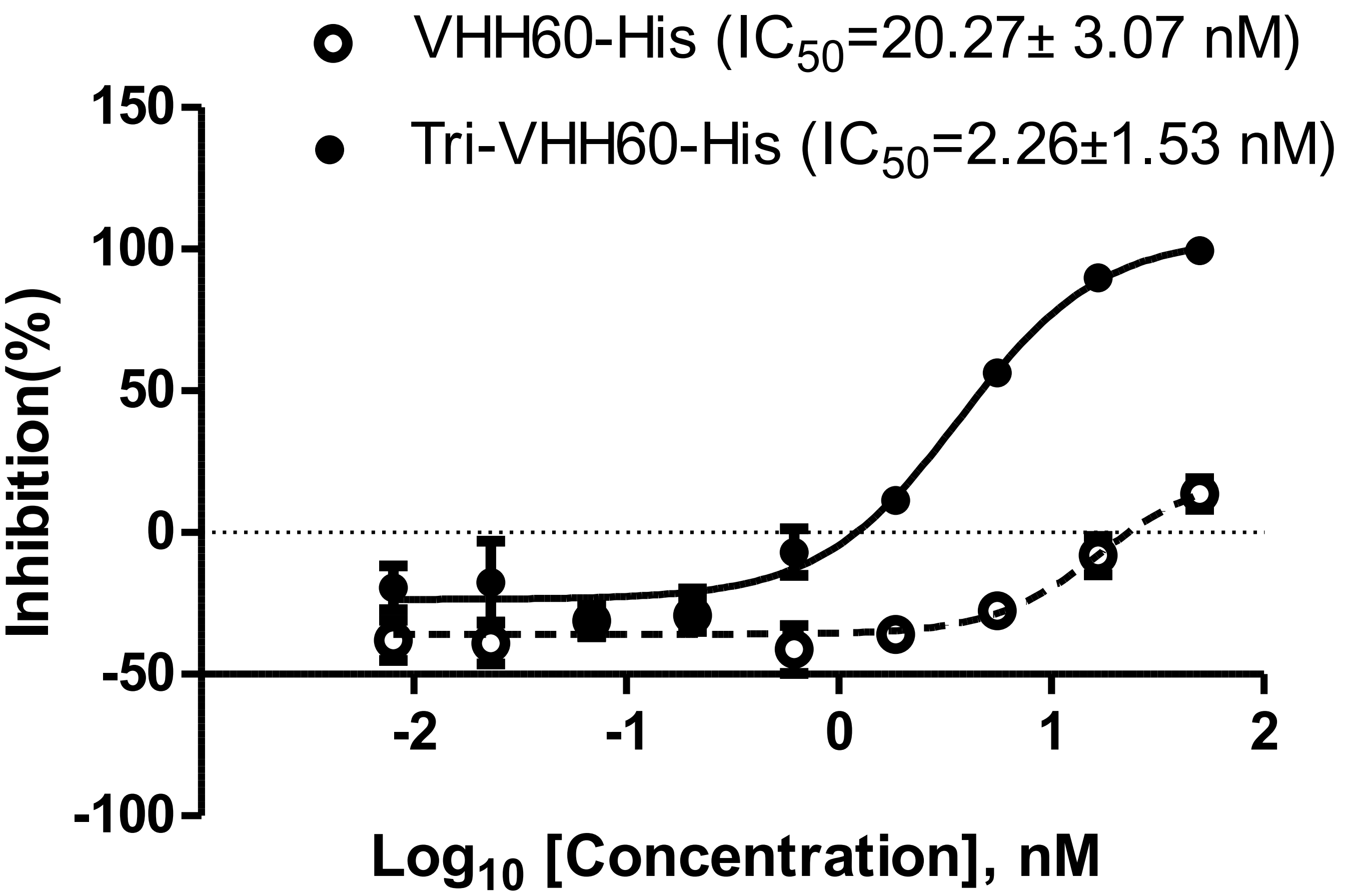

VHH60-His ($IC_{50}$=20.27± 3.07 nM)
Tri-VHH60-His ($IC_{50}$=2.26±1.53 nM)
150
100
50
0
-50
-100
Inhibition(%)
-2
-1
0
1
2
$Log_{10}$ [Concentration], nM
